# Assessing the impact of gamma irradiation on key biological traits of peach fruit fly, *Bactrocera zonata* (Diptera: Tephritidae) under laboratory conditions

**DOI:** 10.64898/2026.03.14.711761

**Authors:** Bibi Hajra, Muhammad Hamayoon Khan, Farrah Zaidi, Muhammad Salman, Zainab Saeed, Usman Khalique, Mehreen Ayaz, Syeda Hira Fatima, Syed Jawad Ahmad Shah

## Abstract

*Bactrocera zonata* (Saunders) (Diptera: Tephritidae) poses a significant threat to global fruit production due to its high reproductive capacity and broad host range. This study aimed to evaluate the effects of gamma irradiation on key biological parameters of *B. zonata*. Six-day-old pupae were exposed to irradiation doses of 0, 30, 40, 50, 60, and 70 Gy, and subsequent developmental and reproductive traits were assessed to determine the impact of irradiation. Post-irradiation results revealed a dose-dependent trend. Higher doses (≥50 Gy) significantly reduced adult emergence, increased the incidence of partially emerged or deformed adults, and shortened adult longevity. Reproductive potential was significantly impaired in males irradiated at 60 and 70 Gy when mated with un-irradiated females, resulting in a marked decline in both fecundity and egg hatchability. Females irradiated at doses ≥50 Gy failed to produce eggs when paired with either irradiated or non-irradiated males, indicating a high level of radio-sensitivity in female flies. Additionally, several traits in the F1 generation such as pupal recovery, pupal size, and adult development exhibited significant abnormalities and suggesting that the effects of irradiation may be transmitted to the next generation. Sterility was highest in males irradiated at 60 and 70 Gy, while females exhibited complete sterility at doses exceeding 40 Gy. The findings indicate that a dose of 70 Gy may be optimal for effective sterility induction in *B. zonata*. However, further detailed studies are required to standardize this dose, incorporating rigorous quality control measures to optimize its application in sterile insect technique (SIT) programs.

## Introduction

Fruit flies (Diptera: Tephritidae) cause significant economic threats to global agriculture, infesting a variety of vegetables and fruits (Khan et al. 2021, Saeed et al. 2022). They deposit their eggs within fruits and vegetables with the help of a pointed, sharp ovipositor. After hatching, the larvae feed on the pulp of the fruit, thereby reducing the quality and quantity of crops (Rattanapum et al. 2009, Javed et al. 2015). Fruit flies can damage crop production ranging from 20% to 80%, but in cases of severe infestation, the entire crop may be perished (Rauf et al. 2013, Khan et al. 2021). Fruit flies are significant quarantine pests, and are often transported along with fruits and vegetables through international trade, and hence have a high potential for spreading and infesting new regions (Khan et al. 2021).

Among the fruit flies, the genus *Bactrocera* of Tephritidae has 651 described species, with approximately 50 of these are significant polyphagous pests (Vargas et al. 2015, Saeed et al. 2022). Pakistan has recorded eleven species of fruit flies, including some of the most dangerous species in the genus *Bactrocera* (Gul et al. 2015, Sarwar et al. 2023). In Pakistan, *B. zonata* (Saunders) (Diptera: Tephritidae), is a serious fruit pest, causing significant economic losses annually. The specie is widely distributed in all provinces of the country (Sarwar et al. 2013, Khan and Naveed 2017, Khan and Akram 2018). The invasive nature of *B. zonata* is attributed to its flight potential, allowing it to travel a distance of at least 25 miles in search of hosts. As a result, its presence and distribution are closely linked with the availability of favorable environment and a suitable host. Additionally, its short generation time and wide-host range allows it to adapt quickly and thrive in a wide range of environments and habitats (Zingore et al. 2020). The presence of host and conducive climatic variables, especially average temperature, plays a vital in determining population density of *B. zonata* (Ghanim and Kontsedalov 2009).

In Pakistan, *B. zonata* incurs an annual economic loss of $200 million to fruit and vegetable growers, with additional costs to other stakeholders involved in the supply chain, including dealers, retailers, and exporters (Rauf et al. 2013). Infestation levels can reach to alarming level of 50% in summer guavas in Pakistan, which results in undersized, deformed and decaying fruits at harvest (Awad et al. 2014). The flies are usually managed by broad-spectrum insecticides. Although effective, the use of insecticides is not environmentally feasible, and also poses risks to non-target organisms and international trade due to toxic residues in fruits (El-Aw et al. 2008) and vegetables. This scenario underscores the importance of increased demand for pest control approaches that are efficient and eco-friendly, such as Sterile Insect Technique (SIT). SIT is a species-specific technique that reduces reliance on chemical pesticides, thereby suppressing pest populations without harming beneficial insects (Vreysen et al.2006).

SIT is a pest management approach involving mass production and release of sterile individuals of the target pests into the environment for mating their wild partners to reduce reproductive potential and minimize their population (Knipling 1955). This technique is safe, environment-friendly and highly target-specific for pest control and is being practiced since 1950s, with no reported risks (Mahmoud and Barta 2011). The application of SIT by the Food and Agriculture Organization/ International Atomic Energy Agency, Vienna Austria (**FAO/IAEA)** remained successful in suppressing or eradicating fruit fly from many countries around the world including Argentina, Chile, Peru, Mexico, Israel, South Africa, Spain, Australia, Guatemala and USA (Hendrichs 2000, Dunn and Follett 2017). Gamma irradiation has high penetration capacity and is a commonly used nuclear technique in SIT programs. It involves exposing insect pupae to a specific dose of radiation for inducing sterility in the targeted pest populations (Bakri et al. 2005). The SIT success depends on the production of high-quality sterile male insects which can effectively compete with wild males and successfully mate with wild females and result in no offspring. Standardization of irradiation doses are crucial to determine the optimal gamma dose that maximizes male competitiveness while reducing somatic damage (Sayed et al. 2013). Therefore, evaluating the effects of radiation on quality control parameters of laboratory-reared fruit flies are essential for successful SIT implementation (Draz et al. 2008, Mahmoud and Barta 2011).

The present study investigated the impact of different gamma irradiation doses on *B. zonata* under controlled laboratory conditions. The primary objectives were to evaluate how irradiation influences key biological parameters and to assess its effectiveness in inducing sterility. The findings may help enhance pest management strategies, particularly those involving SIT.

## Materials and Methods

### Study area

The present study was conducted in the fruit fly rearing laboratory, Plant Protection Division, Nuclear Institute for Food and Agriculture (NIFA), Peshawar, Pakistan. Further, the morphological deformities were examined under a microscope at the Vector Biology Laboratory of the Institute of Zoological Sciences, University of Peshawar, Pakistan.

### Survey of *B. zonata* prevalence in District Peshawar (KP)

Peshawar, the capital of Khyber Pakhtunkhwa Province in Pakistan, is situated at an elevation of 350 meters (Coordinate: 34.0083° N, 71.5189° E) above sea level. The district’s relatively moderate altitude makes it a favorable region for raising variety of crops, including both temperate and subtropical fruits. Peshawar’s climate features hot summers and mild winters, with most of the rainfall occur in the monsoon. The alluvial fertile soil of Peshawar supported by the irrigation system fed by the Kabul River and its tributaries, reinforces the cultivation of various crops. Fruit cultivation in the area includes citrus, mangoes, guavas, and pomegranates, among others. However, similar to other agricultural areas, Peshawar also faces the persistent challenge of fruit fly infestations (see Fig. 1). Therefore, a small survey was conducted to assess the prevalence of *B. zonata* in 32 selected localities within Peshawar district.

**Fig. 1.**
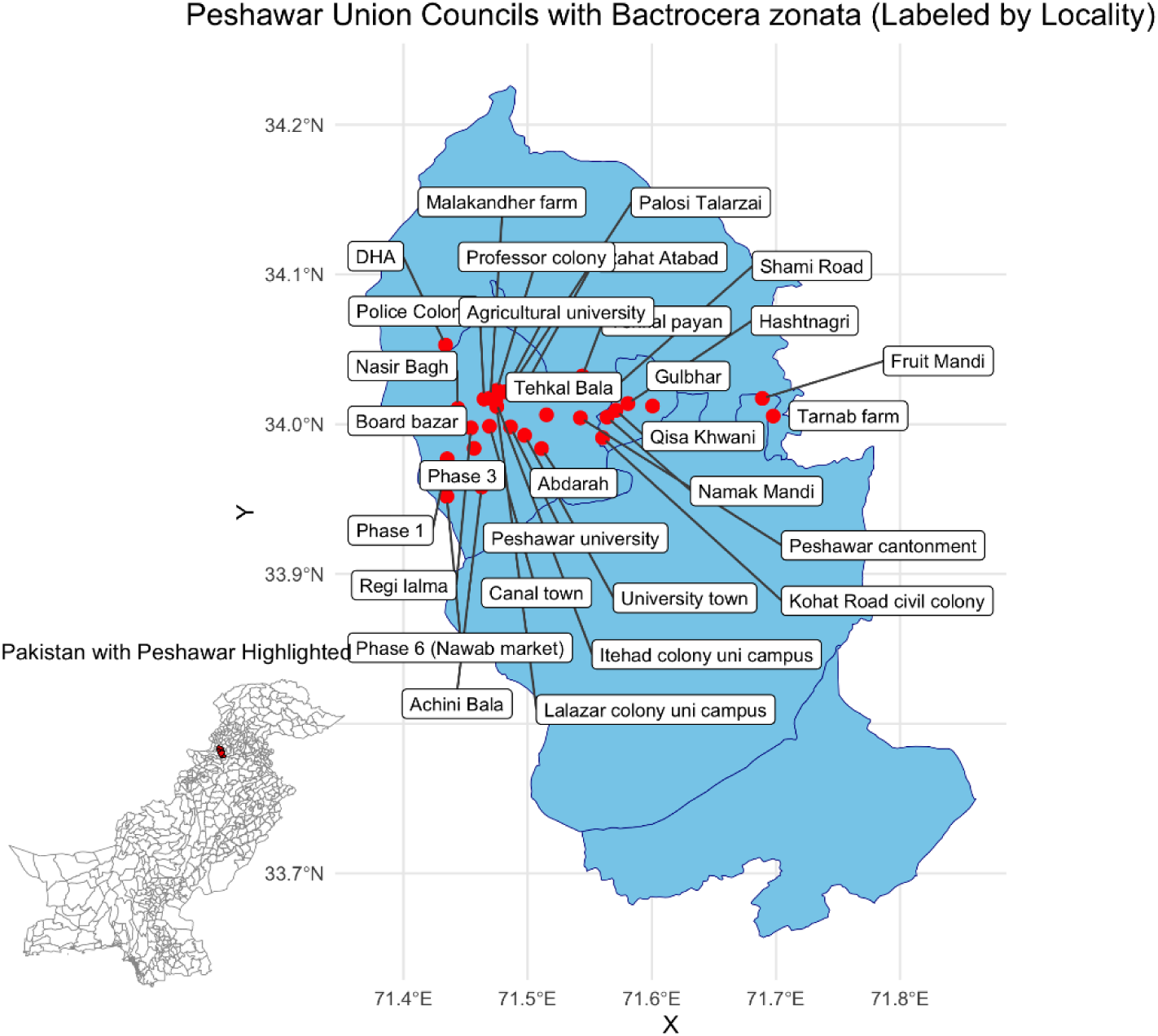
Map of district Peshawar showing the prevalence of *B. zonata* across different Union Councils. The X-axis represents longitudinal coordinates, while the Y-axis represents latitudinal coordinates.

### Rearing of *B. zonata*

Infested fruits samples were collected from various orchards in Peshawar, KP. Trays containing sampled fruits were kept in chamber under controlled conditions (27 ± 1°C, 60 ± 5%) for subsequent pupal formation. Third instar larvae exited the infested fruits and burrowed into sawdust to pupate. Sawdust was examined by sieving after three days and the freshly formed pupae were collected and transferred to adult rearing cages (35 × 30 × 35 cm). After emergence, *B. zonata* adults were provided with a 3:1 mixture of protein hydrolysate, sugar and yeast in small Petri dishes as a food source. A cotton swab partially immersed in a 250 ml glass jar filled with water was supplied to meet the flies hydration needs. Fruits such as bananas and guava were provided to facilitate female oviposition and were regularly replaced with fresh fruit.

### Gamma Irradiation Treatment

Six hundred pupae per treatment group (n=600), approximately 6 days old and close to emergence, were transferred to separate petri dishes. These were then exposed to gamma radiation doses of 30, 40, 50, 60, and 70 Gy using a Co^60^ ISOGAMMALLTYPE high dose irradiator at the Food and Nutrition Division, NIFA, Peshawar. Gamma irradiation was performed at a dose rate of 370 Krad/hr (3.70 KGy/hr). A comparable group of pupae (n=600), also nearing emergence, was maintained as the control group for comparisons. Each does was applied in six replicates, with 100 pupae per replicate (6 × 100 pupae). During irradiation, each replicate was placed in a well-ventilated container, with an alanine reference dosimeter attached to the irradiation chamber to measure the total dose delivered to the pupae. The irradiated replicates were individually maintained in rearing cages to assess various biological parameters of *B. zonata*, including adult emergence, partial emergence, deformities, sex ratio, and adult longevity (in days). The same biological parameters were also recorded for the Control group.

### Effect of Gamma Irradiation on the Biological Parameters of *B. zonata*

#### Adult Emergence

Pupae exposed to the respective irradiation doses were individually transferred to adult fruit fly rearing cages to monitor adult emergence.

The percentage of adult emergence (% Emergence) was calculated using the formula descried by Collins et al. (2008):

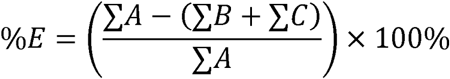

∑ A = number of pupae
∑ B = number of not emerged
∑ C = number of partially emerged

To determine the sex ratio, a cohort of 100 fruit fly pupae was placed in a Petri dish. After a 10-day development period, the emerged adults were counted and classified by sex. The sex ratio was calculated using the formula: number of males/total number of adults.

Adult longevity was assessed by confining freshly emerged flies in rearing cages supplied with adult diet and water-soaked cotton swabs. The cages were monitored daily to record mortality. In addition, irradiated flies were transferred to the Vector Biology Laboratory at the Institute of Zoological Sciences, University of Peshawar, for microscopic examination of morphological deformities, with particular focus on the wings and abdomen. Observed deformities were documented and photographed.

#### Fecundity, Egg Hatchability and Sterility tests

Following complete adult emergence, ten pairs of 12-day-old virgin males and females were selected from each irradiation dose and replicated three times (n = 3). These pairs were maintained in separate cages under the following mating combinations: irradiated male × non-irradiated female, irradiated female × non-irradiated male, irradiated male × irradiated female and non-irradiated male × non-irradiated female (control).

For oviposition, plastic cups with 0.5 mm perforations were internally lined with guava essence and placed inside the adult cages. Eggs laid in these cups were carefully collected, and their number was counted to asses female fecundity. The collected eggs were then transferred to a semi-artificial diet composed of mashed banana (18%), yeast extract (18%), distilled water (60%), Nipagen (2.4%), and sodium benzoate (1.6%) to evaluate hatchability. The number of hatched eggs was recorded, and the hatchability percentage was calculated.

Additionally, male and female fertility across the treatment groups was evaluated based on egg hatchability, which was used to calculate the percentage of sterility. Sterility induced by irradiation was determined using the following formula (Tappozada et al. 1966, Zahran et al. 2013).

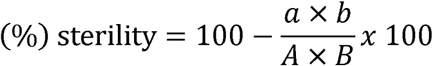

a = No. of eggs laid per female in the treatment.
b = % eggs hatchability in treatment.
A = No. of eggs laid per female in the control.
B = % eggs hatchability in control.

#### Pupal recovery tests

Similarly, data on pupal recovery from each treatment were recorded and analyzed by using the following formula: total number of F1 larvae inoculated/ Number of F1 pupae recovered. Seven-day-old pupae obtained from the mating pairs were used to determine pupal size. To calculate the mean pupal weight (mg), individual pupae were weighed using an electronic balance, and pupal length (cm) was measured with a ruler. Additional biological parameters, including adult emergence, partial emergence, and morphological deformities, were documented.

#### Statistical analysis

All collected data were subjected to statistical analysis to evaluate the effects of gamma irradiation on *B. zonata*. The experiment followed a Completely Randomized Design (CRD) with six radiation dose levels: D0: (0 Gy, Control), D1: 30 Gy, D2: 40 Gy, D3: 50 Gy, D4: 60 Gy, D5: 70 Gy. Each treatment was replicated six times, with 100 pupae per replicate (6 × 100 pupae). The experimental unit for the initial irradiation and emergence studies consisted of a replicate of 100 pupae exposed to a specific irradiation dose.

The biological response variable included adult emergence (%), partial emergence (%), deformities (%), sex ratio (proportion of male), adult longevity (days), fecundity (eggs/female), egg hatchability (%), sterility (%), pupal recovery rate (%), pupal weight (mg), and pupal length (cm).

Subsequent biological parameters, such as the percentage of adult emergence, were calculated using the formula provided by Collins et al. (2008), while partially emerged and deformed adults were recorded directly. The sex ratio was determined using the formula: number of males/total number of adults. Adult longevity was assessed by confining freshly emerged flies in rearing cages.

For assessment of fecundity, egg hatchability, sterility, pupal recovery, and associated F1 parameters, three replicates were conducted for each of the four mating combinations at each irradiation dose. Within each replicate, the experimental unit was a cage containing 10 mating pairs representing one of the following cross combination: irradiated male × non-irradiated female, irradiated female × non-irradiated male, irradiated male × irradiated female and non-irradiated male × non-irradiated female (control). Female fecundity was determined by directly counting the number of eggs laid. These eggs were then transferred to a semi-artificial diet to assess hatchability, and the number of hatched eggs was recorded from the total eggs laid in each replicate. Sterility induced by irradiation was calculated using the formula (Tappozada et al. 1966, Zahran et al. 2013).

Pupal recovery and associated F1 parameters including pupal size, adult emergence, partially emerged adults, and deformities, were assessed from samples of F1 pupae obtained from each replicate of the parental crosses. Pupal size, measured as individual weight (mg) and length(cm), was recorded for a representative sample of pupae from each replicate. Furthermore, the percentage data for various biological parameters of *B. zonata* under different gamma irradiation doses were expressed as means (±SE) of the replicates and analyzed using one-way analysis of variance (ANOVA) in Statistix 8.1 (Analytical Software, Tallahassee, FL). When necessary, data were square root–transformed [√(x + 0.5)] to meet the assumptions of normality and homogeneity of variances, which were tested before analysis. Treatment means were compared using the Least Significant Difference (LSD) test at a significance level of α = 0.05. Untransformed means are reported in all tables and figures.

This statistical approach was employed to provide a comprehensive assessment of the effects of varying gamma irradiation doses on the biological parameters of *B. zonata*.

## Results

### Effect of irradiation doses on the adult emergence of *B. zonata*

Analyses of variance (ANOVA), followed by the Least Significant Difference (LSD) test, were used to statistically compare the biological parameters of *B. zonata* across different irradiation doses. The findings revealed that higher doses (70 Gy and 60 Gy) significantly reduced adult emergence compared to the control group, which exhibited the highest emergence rate (F = 1476.00; df = 5, 30; P < 0.000) (Table 1).

**Table 1.** Effect of different irradiation doses (0, 30, 40, 50, 60, and 70 Gy) on adult emergence (%), partial emergence(%), and % deformity (%) of *B. zonata*. Data are as means ± standard error (SE). Different letters denote statistically significant differences among treatments (P≤0.05, LSD test).

| **Dose**<br>**(Gy)** | **Adult emergence(%)** | **Partial**<br>**emergence(%)** | **Deformity(%)** |
| --- | --- | --- | --- |
| 0 | 95.33 ± 0.88 A | 0.5 ± 0.28 D | 0.17 ± 0.17 B |
| 30 | 85.33 ± 0.72 B | 4.5 ± 0.57 C | 1.75 ± 0.01 B |
| 40 | 83.5 ± 0.86 B | 4.5 ± 1.15 C | 1.81 ± 0.71 B |
| 50 | 38.0 ± 1.15 C | 9.66 ± 0.72 B | 6.49 ± 1.32 B |
| 60 | 37.0 ± 0.28 C | 16.5 ± 0.86 A | 9.42 ± 2.26 B |
| 70 | 29.5 ± 0.28 D | 18.0 ± 0.57 A | 29.24 ± 7.05 A |
| LSD | 2.372 | 2.30 | 9.5 |

Similarly, higher irradiation doses had a pronounced effect on other biological traits, significantly increasing the percentage of partial emergence (F = 89.40; df = 5, 30; P < 0.000) and deformities (F = 12.50; df = 5, 30; P < 0.0002) compared to the control group (0 Gy), which recorded the lowest mean percentages for both paraments (Table 1).

Gamma irradiation at a certain dose level can induce cellular damage in pupae, resulting in various emergence-related abnormalities. For example, at higher doses (50, 60, and 70 Gy), some flies only partially emerged from the pupal casing (Fig. 2A and B), while others fully emerged but were either dead or exhibited physically deformities (Fig. 3A and B). In more severe cases, irradiation completely inhibited pupal emergence at doses of 30, 40, 50, 60, and 70 Gy (Fig. 4).

**Fig. 2.**
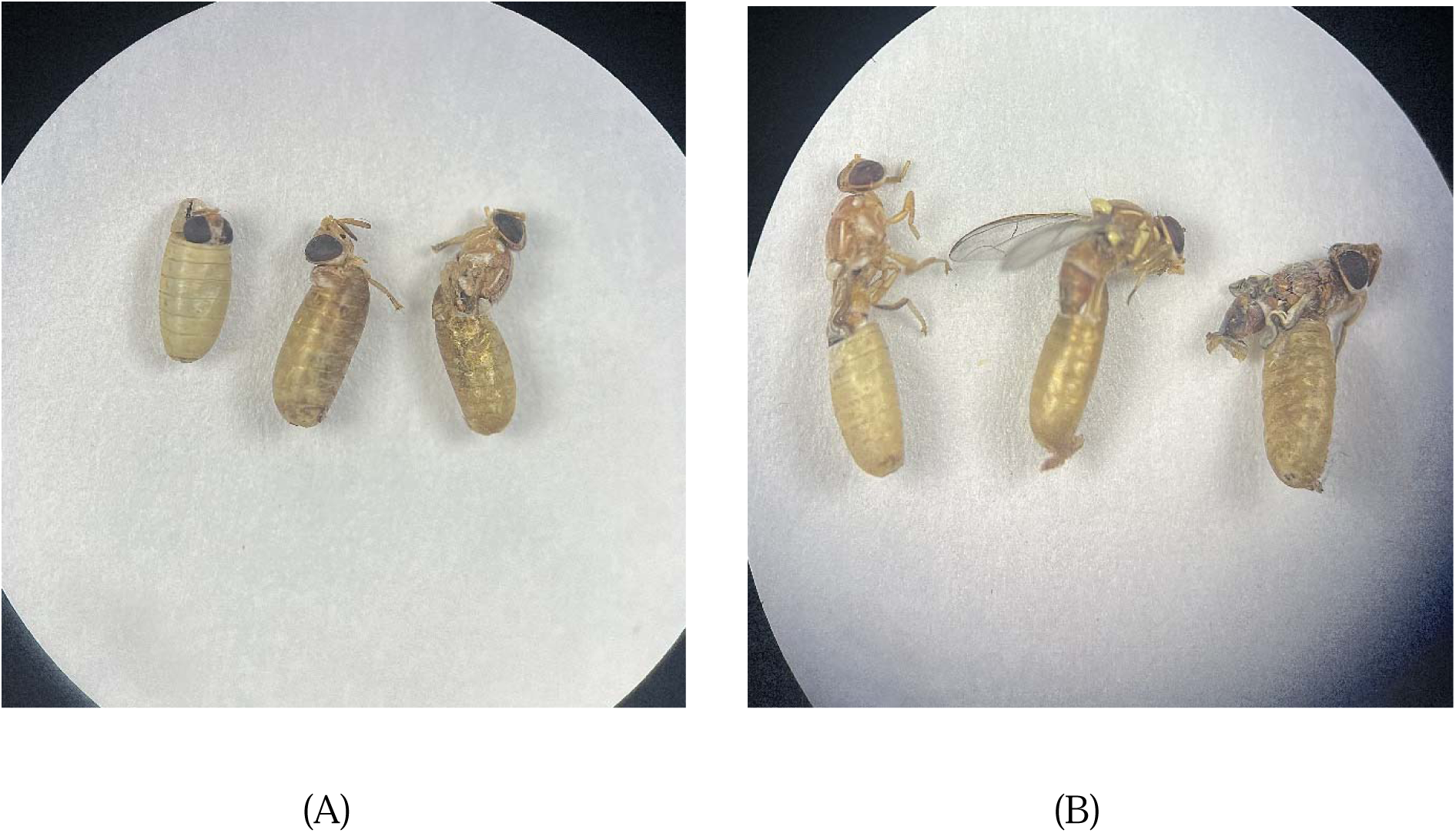
Partially emerged *B. zonata* showing pronounced effects at higher gamma irradiation doses (50, 60, 70 Gy). (A) Pupae exhibiting incomplete adult emergence, with head and thorax visible while the wings and abdomen remain enclosed. (B) Adults displaying partial wing expansion or remaining trapped within the pupal case, with deformed wings.

**Fig. 3.**
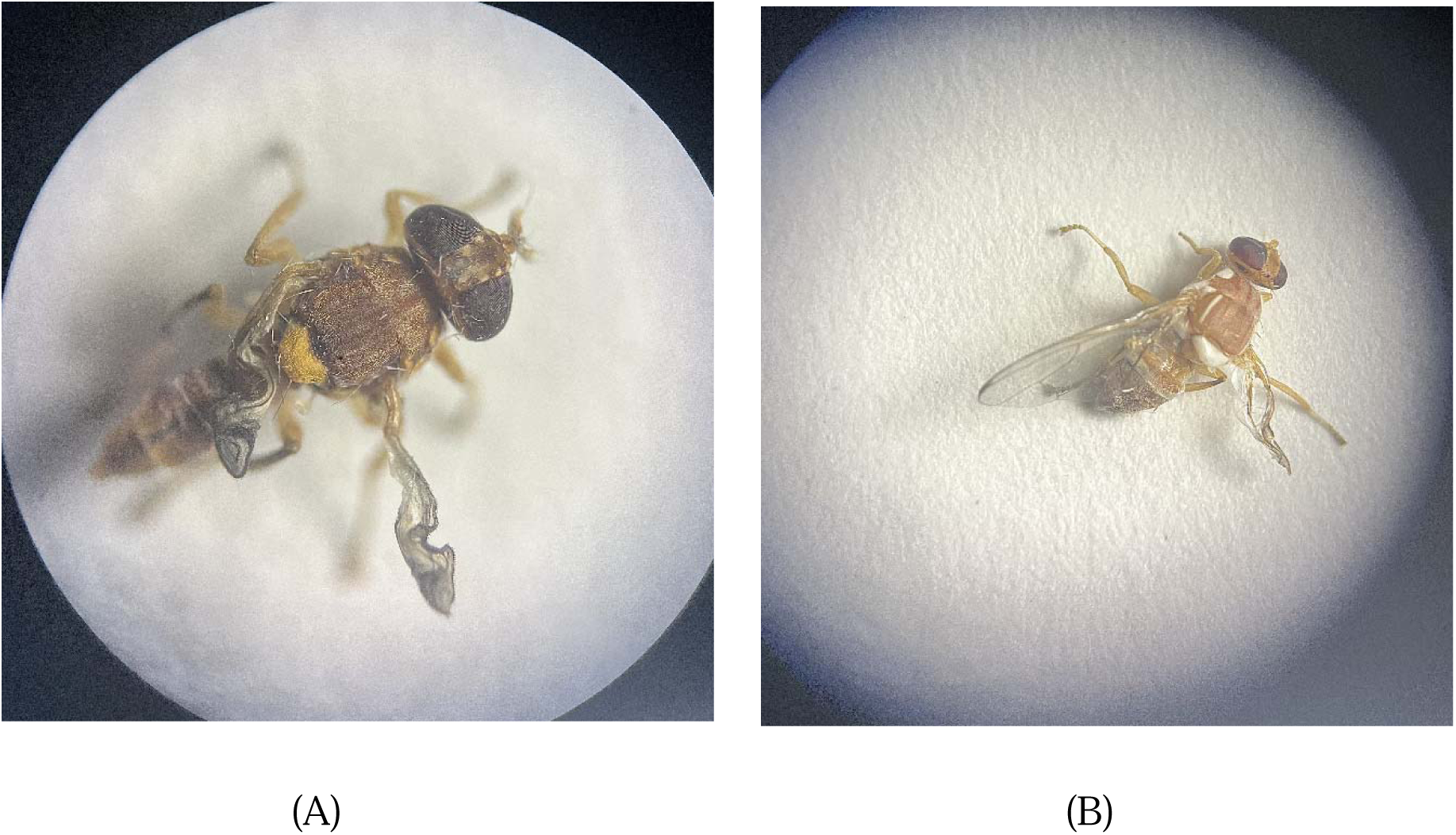
illustration of wing deformities and abdominal constriction in *B. zonata* observed at higher gamma irradiation doses (50, 60, 70 Gy). (A) Adult exhibiting severely crumpled wings and a constricted abdomen, indicating abnormal development. (B) Adult with partially expanded wings showing noticeable malformation and body deformities.

**Fig. 4.**
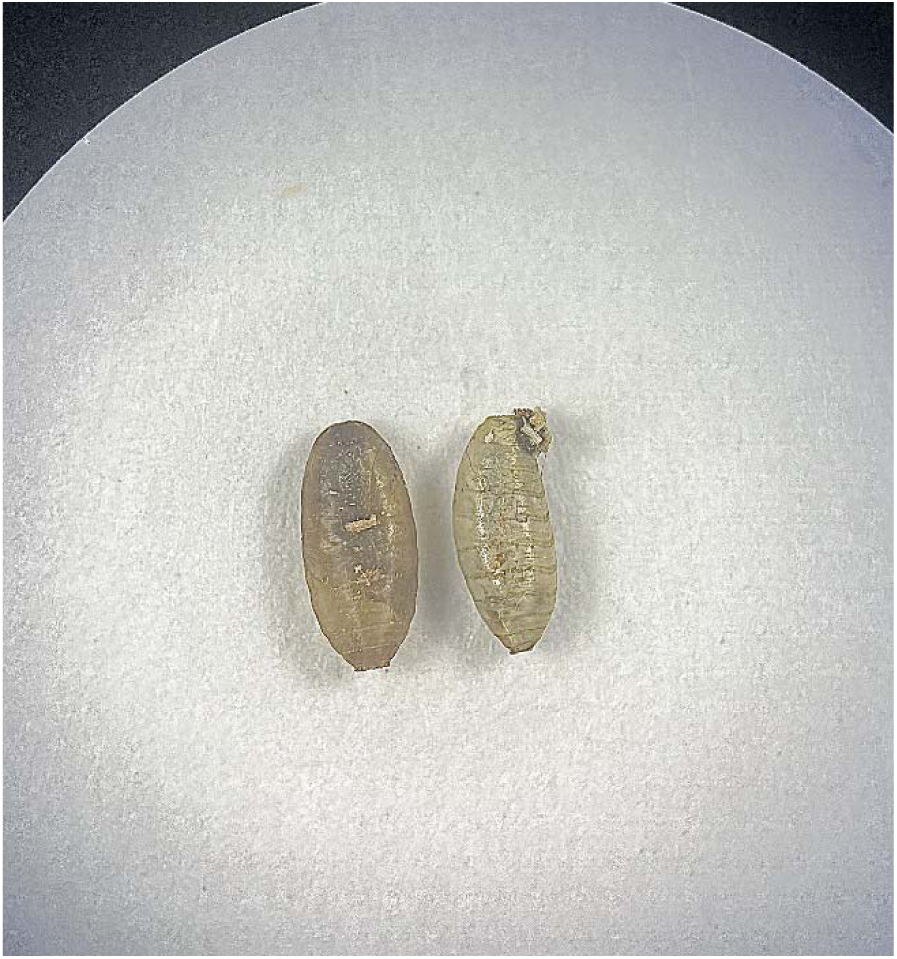
Unhatched pupae of *B. zonata* resulting from exposure to various gamma irradiation doses (30, 40, 50, 60, 70 Gy).

### Effect of irradiation doses on the sex ratio of *B. zonata*

The results indicated that gamma irradiation had a significant effect on the sex ratio of *B. zonata.* The highest proportion of female progeny was observed at 30 Gy, while the lowest was recorded at 70 Gy, followed by 60 Gy and 40 Gy (F = 6.97; df = 5, 30; P < 0.0029) (Figure 5).

**Fig. 5.**
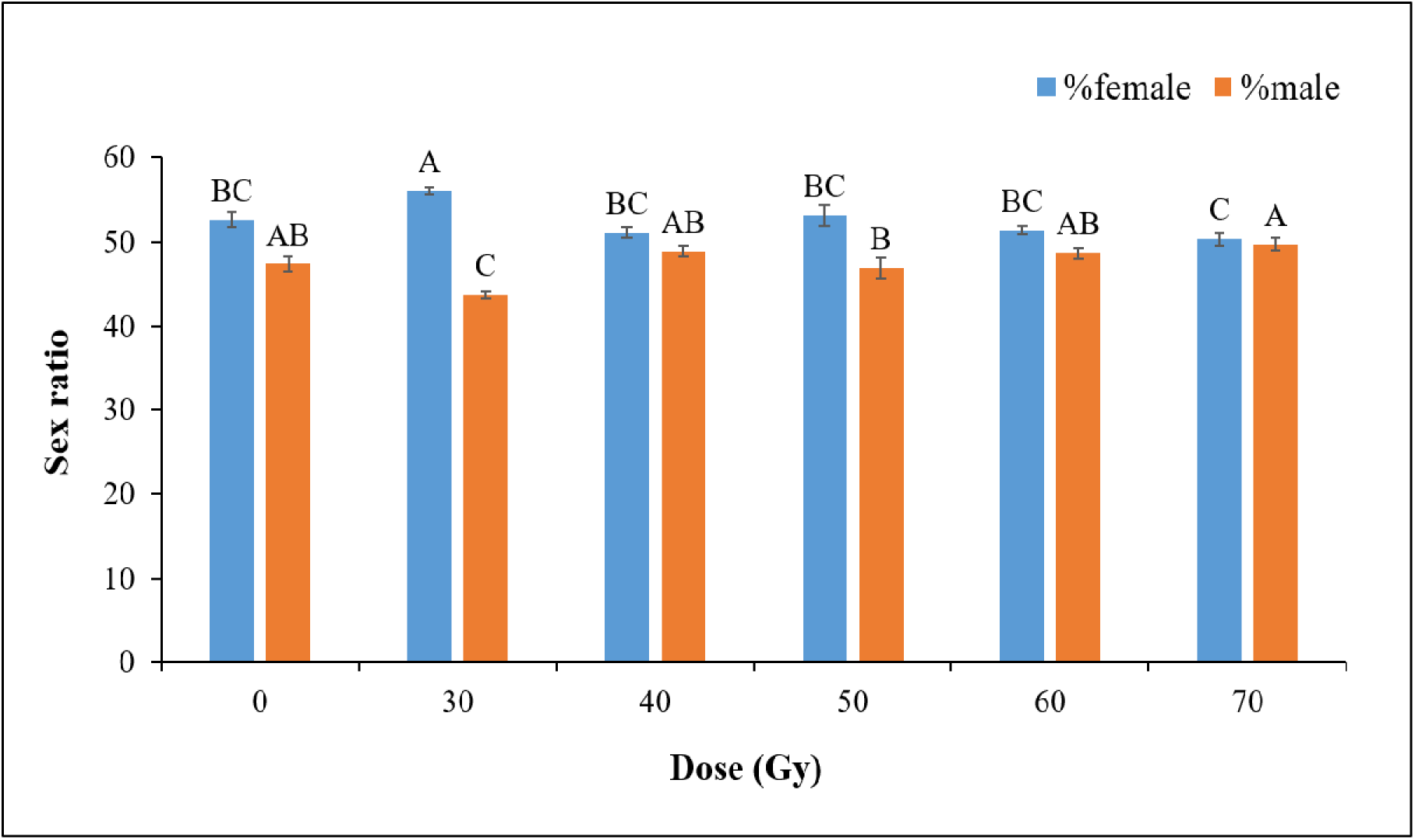
Sex ratio of B. zonata emerging from pupae exposed to different gamma irradiation doses (0, 30, 40, 50, 60, and 70 Gy). The results indicate a slight predominance of female emergence across all treatments. Data are presented as means ± standard error (SE). Different letters indicate statistically significant differences among treatments (P≤0.05, LSD test).

In contrast, the highest percentage of male progeny was recorded at 70 Gy. While the lowest was observed at 30 Gy, followed by 50 Gy (F = 6.62; df = 5, 30; P < 0.0035) (Figure 5).

### Effect of irradiation doses on the longevity (days) of *B. zonata*

The longevity of adult flies emerging from irradiated pupae was significantly affected by the gamma irradiation dose. The longest lifespan in females was observed in the control group, while the shortest was recorded at 70 Gy, followed by 60 Gy and 50 Gy (F = 365.00; df = 5, 30; P < 0.00) (Figure 6).

**Fig. 6.**
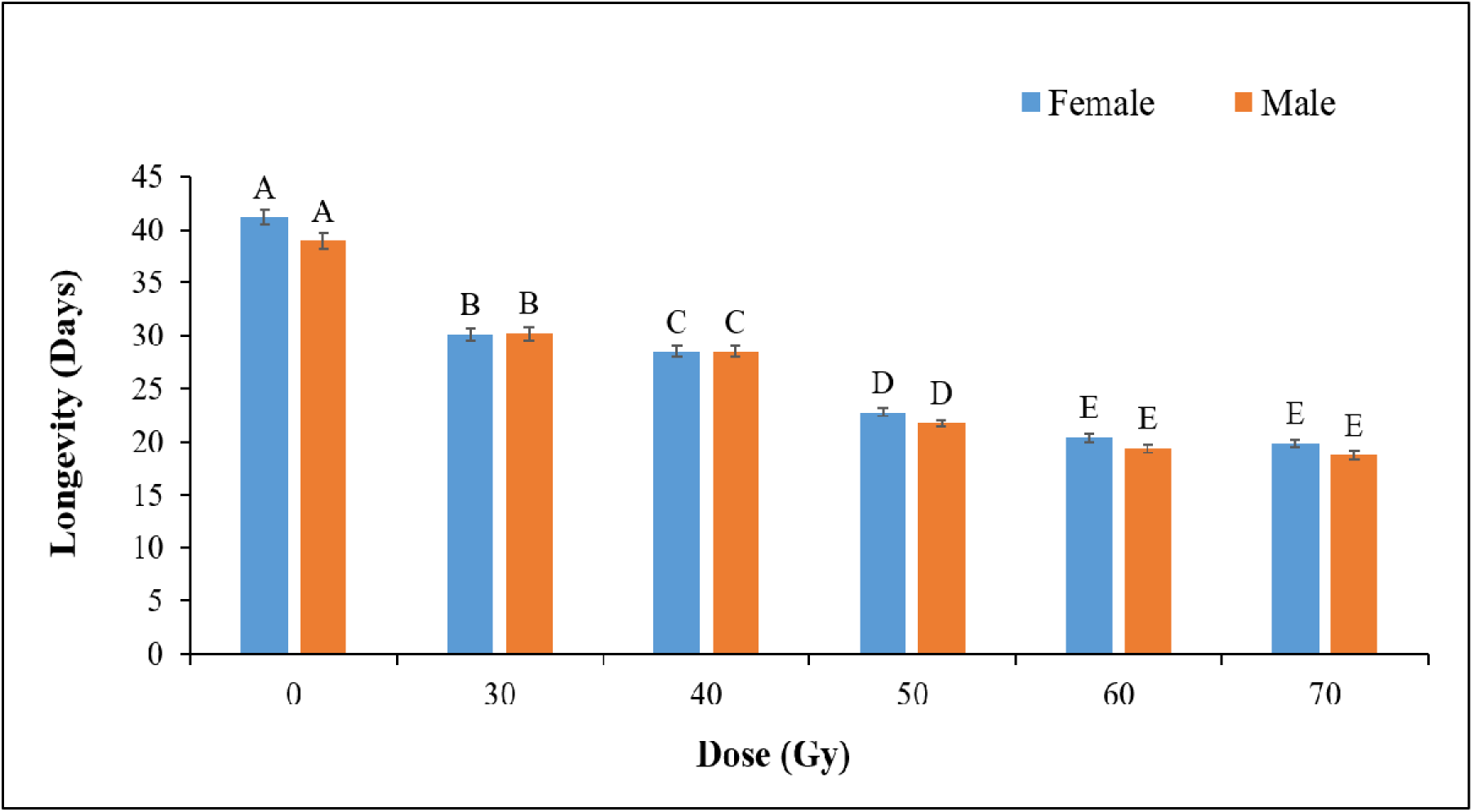
Effect of different gamma irradiation doses (0, 30, 40, 50, 60, and 70 Gy) on the longevity (days) of B. zonata adults. The results show a significant decrease in longevity with increasing irradiation doses, with the shortest lifespan observed at 70 Gy. Data are presented as means ± standard error (SE). Different letters indicate statistically significant differences among treatments (P≤0.05, LSD test).

Similarly, male flies also exhibited the highest longevity in the control group, while the shortest lifespan was recorded at 70 Gy, followed by 60 Gy and 50 Gy (F = 491.00; df = 5, 30; P < 0.000) (Figure 6).

### Effect of irradiation doses on the fecundity and fertility after crossing of irradiated male and un-irradiated female of *B. zonata* (IrM × UnIrF)

The results revealed that different irradiation doses significantly affected the fecundity and fertility of *B. zonata* when irradiated males were mated with un-irradiated females (IrM × UnIrF). The lowest mean fecundity per female was recorded at 70 Gy, followed by 60 Gy, while, the highest fecundity was observed in the control group (F = 470.00; df = 5, 12; P < 0.000). Likewise, the lowest egg hatchability was recorded at 70 Gy, followed by 60 Gy and 50 Gy, whereas, the highest hatchability was observed in the control group (F = 1222.00; df = 5, 12; P < 0.000) (Table 2).

**Table 2.**
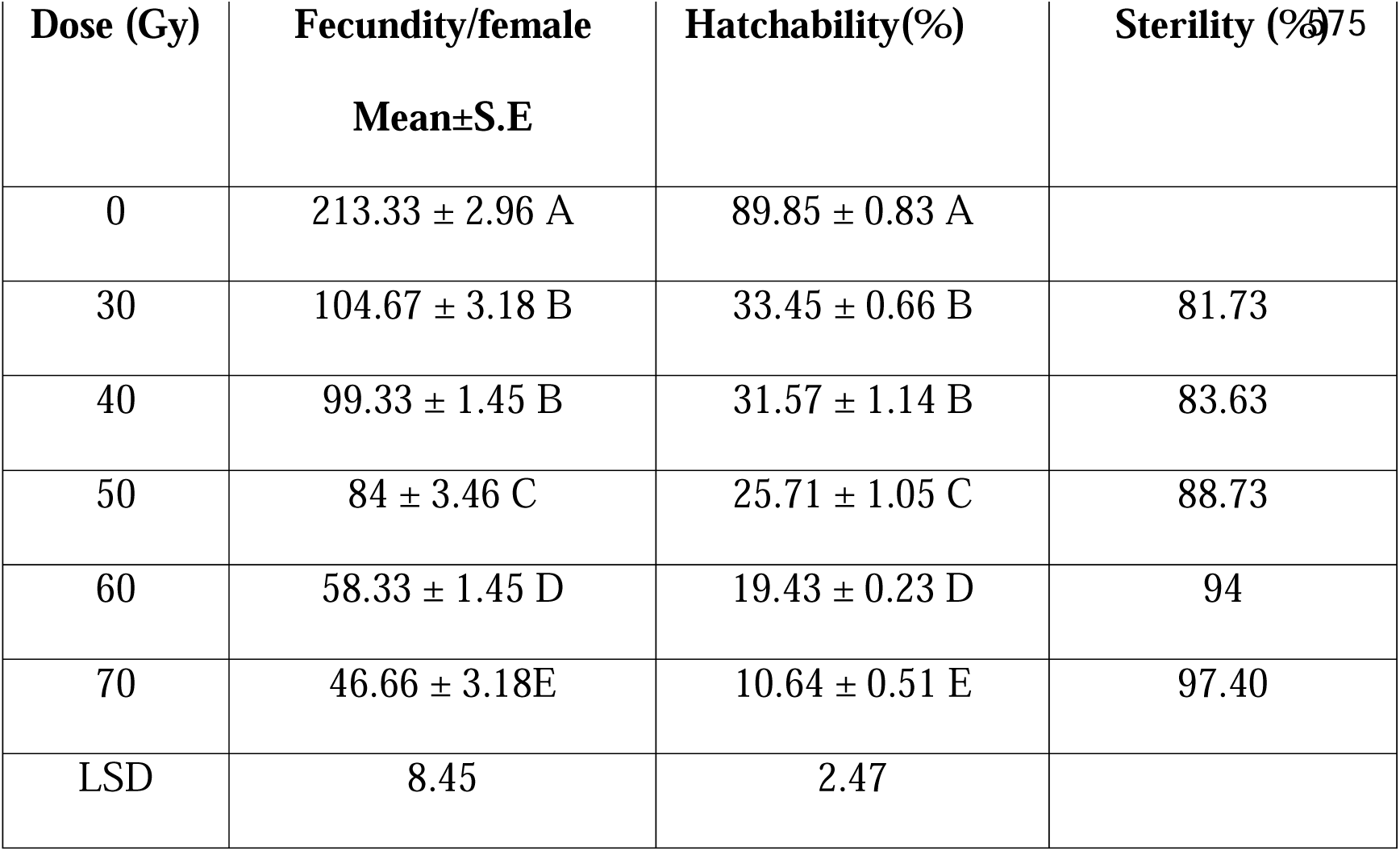
Fecundity per female, hatchability(%), and sterility(%) of *B. zonata* resulting from e crosses between irradiated male and un-irradiated female (IrM+UnIrF). Data are presented as means ± standard error (SE). Different letters indicate statistically significant differences among treatments (P≤0.05, LSD test).

Furthermore, the highest percentage of sterility was observed at 70 Gy, followed by 60 Gy and 50 Gy (Table 2).

### Effect of irradiation doses on the biological parameters of F1 progeny from crossing of irradiated male and un-irradiated female of *B. zonata* (IrM × UnIrF)

ANOVA followed by the LSD test was used to statistically compare the biological parameters of F1 progeny resulting from the mating of irradiated males and un-irradiated females of *B. zonata* (Table 3). Gamma irradiation had a significant effect on pupal recovery, pupal weight, pupal length, percentage of adult emergence, partial emergence, and deformity.

**Table 3:** Effect of different gamma irradiation doses on pupal recovery, pupal weight, pupal length, adult emergence(%), partial emergence(%), and deformity (%) in *B. zonata* resulting from crosses between irradiated males and un-irradiated females (IrM+UnIrF). Data are presented as means ± standard error (SE). Different letters indicate statistically significant differences among treatments (P≤0.05, LSD test).

| **Dose**<br>**(Gy)** | **Pupal**<br>**recovery** | **Pupal**<br>**weight**<br>**(mg)** | **Pupal**<br>**length**<br>**(cm)** | **Adult**<br>**emergence(**<br>**%)** | **Partial**<br>**emergence(**<br>**%)d** | **%**<br>**Deformity(**<br>**%)** |
| --- | --- | --- | --- | --- | --- | --- |
| 0 | 423 ± 0.57<br>A | 12.9 ± 0.70<br>A | 0.5 ± 0.01 A | 96.23 ± 0.49<br>A | 0 ± 0 D | 0 |
| 30 | 123 ± 1.73<br>B | 11.7 ± 0.18<br>AB | 0.48 ± 0 A | 78.04 ± 0.30<br>B | 0 ± 0 D | 0 |
| 40 | 119 ± 0.57<br>B | 11.4 ± 2.54<br>AB | 0.48 ± 0.01<br>A | 73.94 ± 0.61<br>C | 0.84 ± 0.48 D | 0 |
| 50 | 69 ± 2.30<br>C | 11.4 ± 2.54<br>AB | 0.44 ± 0.02<br>B | 31.84 ± 0.60<br>D | 7.32 ± 1.08 C | 0 |
| 60 | 65 ± 1.73<br>C | 10.1 ± 0.06<br>AB | 0.4 ± 0 C | 30.62 ± 2.74<br>D | 10.88 ± 2.07<br>B | 0 |
| 70 | 47 ± 1.76<br>D | 7.33 ± 0.88 C | 0.38 ± 0.01<br>C | 26.76 ± 0.90<br>E | 25.39 ± 1.29<br>A | 0 |
| LSD | 4.89 | 3.51 | 0.37 | 3.85 | 3.41 | 0 |

The lowest pupal recovery was recorded at 70 Gy, followed by 60 Gy and 50 Gy, whereas the highest mean pupal recovery was observed in the control group (F = 5136.00; df = 5, 12; P < 0.000). Pupal weight was highest in the control group and lowest at 70 Gy (F = 2.68; df = 5, 12; P < 0.0754). Similarly, the shortest pupae were observed at 70Gy, while the longest were recorded in the control group (F = 17.10; df = 5, 12; P < 0.000) (Table 3).

The lowest percentage of adult emergence was recorded at 70 Gy, while the highest was observed in the control group (F = 64.20; df = 5, 12; P < 0.000). The highest proportion of partially emerged adults occurred at 70 Gy, whereas the lowest was observed at 40Gy, with no partial emergence recorded in the control group (F = 33.00; df = 5, 12; P < 0.000) (Table 3). No deformed flies were observed in any treatment group.

### Effect of irradiation doses on the Fecundity and fertility after crossing of irradiated female and un-irradiated male of *B. zonata* (IrF × UnIrM)

The result revealed that fecundity and sterility were significantly influenced by different irradiation doses when irradiated females were crossed with un-irradiated males (IrF × UnIrM). The lowest mean fecundity per female was observed at 40 Gy, followed by 30 Gy, while the highest fecundity was recorded in the control group. Notably, females that emerged from pupae exposed to the higher doses of (50, 60 and 70 Gy) did not lay any eggs (F = 1929.00; df = 5, 12; P < 0.000) (Table 4).

**Table 4:** Fecundity per female, hatchability(%), and sterility(%) of *B. zonata* resulting from crosses between irradiated females and un-irradiated males (IrF+UnIrM). Data are presented as means ± standard error (SE). Different letters indicate statistically significant differences among treatments (P≤0.05, LSD test).

| Dose (Gy) | Fecundity/female | Hatchability(%) | Sterility(%)<br>590 |
| --- | --- | --- | --- |
| 0 | 213.33 ± 2.96 A | 89.85 ± 0.83 A | |
| 30 | 26 ± 2.88 B | 33.67 ± 1.85 B | 95.6 |
| 40 | 25.06 ± 2.18 B | 27.70 ± 3.57 C | 96.37 |
| 50 | 0 ± 0 C | 0 ± 0 D | 100 |
| 60 | 0 ± 0 C | 0 ± 0 D | 100 |
| 70 | 0 ± 0 C | 0 ± 0 D | 100 |
| LSD | 5.88 | 5.17 |  |

The lowest egg hatchability of *B. zonata* was recorded at 40 Gy, followed by 30 Gy, while, the highest hatchability was observed in the control group (F = 438.00; df = 5, 12; P < 0.000). Complete (100%) sterility was observed at irradiation doses of 50, 60 and 70 Gy (Table 4).

### Effect of irradiation doses on the biological parameters of F1 progeny from crossing of irradiated female and un-irradiated male of *B. zonata* (IrF × UnIrM)

The results revealed that gamma irradiation significantly affected the biological parameters of F1 progeny resulting from crosses between irradiated females and un-irradiated males (IrF × UnIrM). The lowest mean pupal recovery was recorded at 40 Gy, followed by 30 Gy, while the highest recovery was observed in the control group. No pupal recovery occurred at 50, 60 or 70 Gy due to complete egg sterility and zero hatchability (F = 55356.00; df = 5, 12; P < 0.000) (Table 5).

**Table 5:**
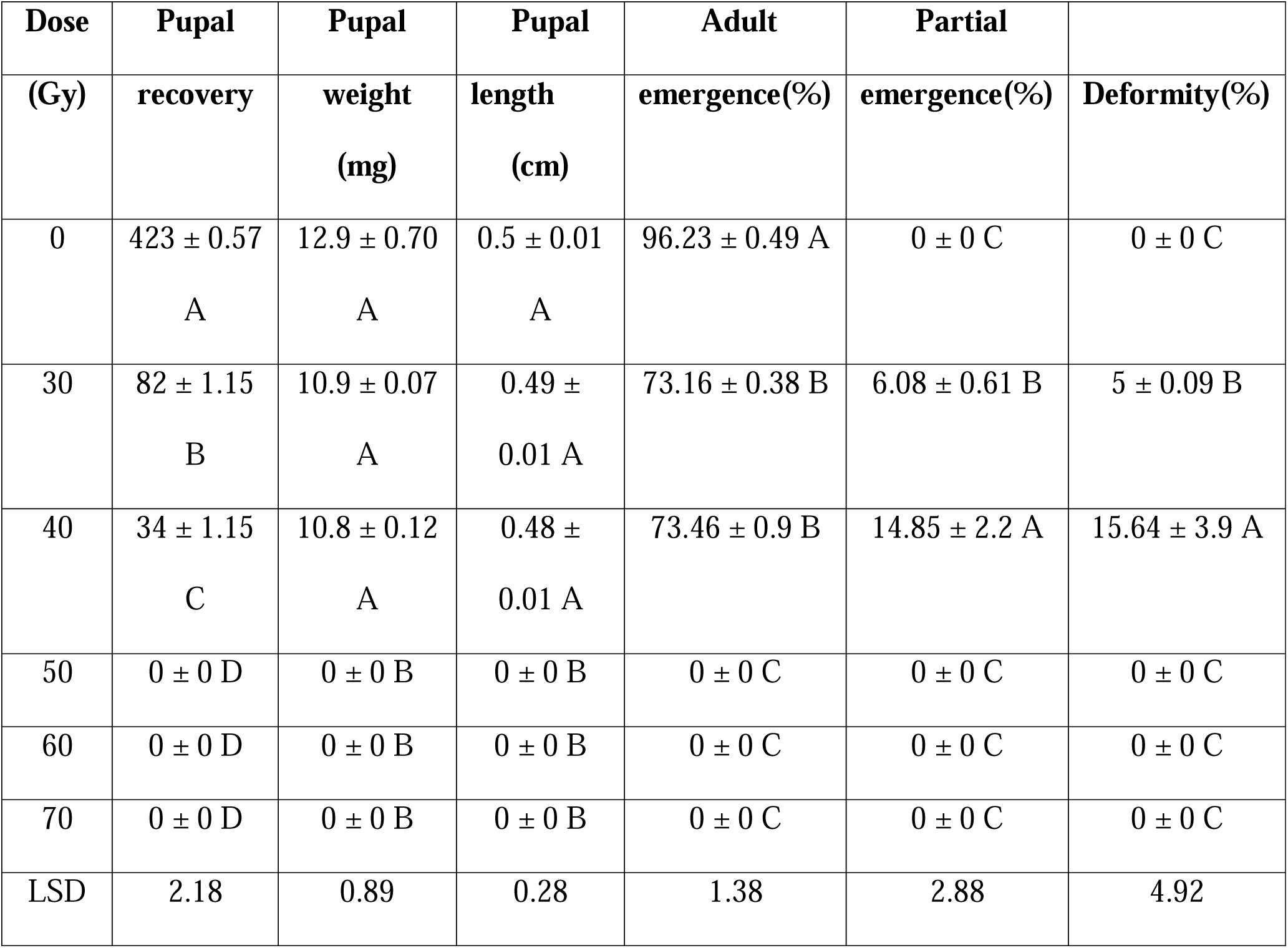
Effect of different gamma irradiation doses on pupal recovery, pupal weight, pupal length, adult emergence(%), partial emergence(%), and deformity(%) in *B. zonata* resulting from crosses between irradiated females and un-irradiated males (IrF+UnIrM). Data are presented as means ± standard error (SE). Different letters indicate statistically significant differences among treatments (P≤0.05, LSD test).

Pupal weight was highest in the control group and lowest at 40 Gy, followed by 30 Gy (F = 436.00; df = 5, 12; P < 0.000). The shortest pupae were observed at 40 Gy and 30 Gy, although the difference between these two doses was not statistically significant. The longest pupae were recorded in control group (F = 864.00; df = 5, 12; P < 0.000) (Table 5).

The lowest percentage of adult emergence was observed at 30 Gy and 40 Gy, while the highest was recorded in the control group (F = 10135.00; df = 5, 12; P < 0.000). The highest percentage of partially emerged flies was observed at 40 Gy, with no partial no partial emergence recoded in the control group (F = 42.20; df = 5, 12; P < 0.000). The mean percentage of deformed flies differed significantly between 40 Gy and 30 Gy, whereas no deformities were observed in the control group (F = 15.60; df = 5, 12; P < 0.0001) (Table 5).

### Effect of irradiation doses on the Fecundity and fertility after crossing of irradiated male and female of *B. zonata* (IrM × IrF)

Significant differences in fecundity and fertility were observed across irradiation doses when both male and female *B. zonata* were irradiated and crossed (IrM × IrF). The lowest mean fecundity per female was recorded at 40 Gy, followed by 30 Gy, while the highest was observed in the control treatment (F = 4011.00; df = 5, 12; P < 0.000) (Table 6).

**Table 6:** Fecundity per female, hatchability(%), and sterility(%) of *B. zonata* resulting from crosses between irradiated males and females (IrM+IrF). Data are presented as means ± standard error (SE). Different letters indicate statistically significant differences among treatments (P≤0.05, LSD test).

| Dose (Gy) | Fecundity/female<br>Mean±S.E | Hatchability(%) | Sterility(%) |
| --- | --- | --- | --- |
| 0 | 213.33 ± 2.96 A | 89.85 ± 0.83 A |  |
| 30 | 28 ± 1.15 B | 20.22 ± 0.64 B | 97.04 |
| 40 | 20.67 ± 0.67 C | 17.73 ± 1.46 C | 98.08 |
| 50 | 0 ± 0 D | 0 ± 0 D | 100 |
| 60 | 0 ± 0 D | 0 ± 0 D | 100 |
| 70 | 0 ± 0 D | 0 ± 0 D | 100 |
| LSD | 4.08 | 2.26 |  |

No egg hatchability was observed from females exposed to 50, 60, or 70 Gy, indicating complete sterility at these doses. In contrast, the highest hatchability was recorded in the control group (F = 4327.00; df = 5, 12; P < 0.000) (Table 6). Complete (100%) sterility occurred at 50, 60 and 70 Gy doses (Table 6).

### Effect of irradiation doses on the biological parameters of F1 progeny from crossing of irradiated male and female of *B. zonata* (IrM × IrF)

Various irradiation doses had a significant, dose-dependent effect on pupal recovery, pupal weight, pupal length, percentage of adult emergence, partial emergence, and deformities in F1 progeny of *B. zonata* resulting from crosses between irradiated males and females (IrM × IrF)

The lowest mean pupal recovery was recorded at 40 Gy, followed by 30 Gy, while the highest recovery was observed in the control group. No pupal recovery occurred at 50, 60, and 70 Gy due to zero egg hatchability in females that developed from irradiated pupae at these higher doses (F = 86616.00; df = 5, 12; P < 0.000) (Table 7).

**Table 7:** Effect of different gamma irradiation doses on pupal recovery, pupal weight, pupal length, adult emergence(%), partial emergence(%), and deformity(%) in *B. zonata* resulting from crosses between irradiated males and females (IrM+IrF). Data are presented as means ± standard error (SE).Different letters indicate statistically significant differences among treatments (P≤0.05, LSD test).

| **Dose**<br>**(Gy)** | **Pupal**<br>**recovery** | **Pupal**<br>**weight**<br>**(mg)** | **Pupal**<br>**length**<br>**(cm)** | **Adult**<br>**emergence(%)** | **Partial**<br>**emergence(%)** | **Deformity(%)** |
| --- | --- | --- | --- | --- | --- | --- |
| 0 | 423 ±<br>0.57 A | 12.9 ±<br>0.70 A | 0.5 ± 0.01<br>A | 96.22 ± 0.49 A | 0 | 0 |
| 30 | 21 ± 0.57<br>B | 10.7 ±<br>0.07 B | 0.46 ± 0<br>A | 84.09 ± 3.23 B | 0 | 0 |
| 40 | 16 ± 1.15<br>C | 10.7 ±<br>0.07 B | 0.42 ±<br>0.02 B | 74.97 ± 3.15 C | 0 | 0 |
| 50 | $0 \pm 0 D$ | $0 \pm 0 C$ | $0 \pm 0 C$ | $0 \pm 0 D$ | 0 | 0 |
| 60 | $0 \pm 0 D$ | $0 \pm 0 C$ | $0 \pm 0 C$ | $0 \pm 0 D$ | 0 | 0 |
| 70 | $0 \pm 0 D$ | $0 \pm 0 C$ | $0 \pm 0 C$ | $0 \pm 0 D$ | 0 | 0 |
| LSD | 1.78 | 0.89 | 0.03 | 5.71 | 0 | 0 |

Pupal weight of *B. zonata* was the highest in the control group and lowest at 30 Gy and 40 Gy, although the difference was not statistically significant (F = 433.00; df = 5, 12; P < 0.000) (Table 7). Similarly, the shortest pupae were recorded at 40 Gy, followed by 30 Gy, while the longest pupae were observed in the control group (F = 651.00; df = 5, 12; P < 0.000) (Table 7).

The lowest percentage of adult emergence was observed in 40 Gy, while the highest was recorded in the control group (F = 645.00; df = 5, 12; P < 0.000) (Table 7). No partially emerged or deformed flies were observed in any of the treatments.

## Discussion

The present study evaluated the effects of various gamma irradiation doses (30, 40, 50, 60, and 70 Gy) on key biological parameters of the peach fruit fly *B. zonata*. The findings indicated that doses of 60 and 70 Gy were the most effective, significantly impacting critical life history and reproductive traits. These higher doses resulted in the lowest adult emergence rates and the highest frequencies of partial emergence and deformities. Such outcomes suggest that high radiation levels can induce physiological damage, leading to reduced numbers of healthy adult flies (Akman and Zumreoglu 1978, Draz et al. 2008, Panduranga et al. 2021). Similarly, Nasution et al. (2018) reported a 66% adult emergence of *B. dorsalis* at 110 Gy.

In our study, the percentage of deformed adults was 9.42% at 60 and 29.24% at 70 Gy. These results are partially consistent with the findings of Zahran et al. (2013), who reported adult malformation rates of 8.3% and 11.7% in *B. zonata* at 70 and 90 Gy, respectively. The observed deformities are likely due to internal developmental disruptions caused by irradiation, leading to physical malformation such as shrunken bodies or wrinkled wings (Nasution et al. 2018, Panduranga et al. 2021).

The sex ratio of emerged adults was slightly skewed towards females at all irradiation doses, with the highest percentage observed at 30 Gy (56.05±0.39%) and the lowest at 70 Gy (50.3±0.75%). In contrast, Zahran et al. (2013) reported an increase in the proportion of females with radiation dose. Conversely, Nasution et al, (2018) observed a decline in the female percentage at higher doses (90 and 110 Gy).

Adult longevity was also significantly affected by irradiation. Aside from the control group, the longest lifespan was recorded at 30 Gy, with progressively shorter lifespans observed at higher doses. Similar findings were reported by Panduranga et al. (2021), who observed reduced longevity in *B. cucurbitae* with increasing irradiation doses. However, Puanmanee et al. (2010) found no significant effect on the longevity of *B. correcta*, likely due to the lower doses used (0, 5, 10, and 30 Gy) and differences in species-specific responses.

Fecundity decreased significantly with increasing irradiation doses, likely due to the effects of ionizing radiation on living cells. This process involves a cascade of oxidative reactions along the radiation path, resulting in the formation of harmful free radicals, particularly peroxy radicals which can damage cellular structures and impair reproductive function. These free radicals cause irreversible damage to organic molecules, leading to chromosome fragmentation and various chromosomal aberrations, including dominant lethal mutations and translocations. Such disruption result in the production of imbalanced gametes, and inhibit mitosis, often causing the death of fertilized eggs. As dividing cells, particularly germ cells, are highly sensitive to radiation, reproductive tissues are especially vulnerable. (Expo et al. 2015). The lowest fecundity per female (46.66%), was observed when a normal female was mated with a male irradiated at 70 Gy. Complete sterility (zero fecundity) was recorded at 50, 60, and 70 Gy when irradiated females were crossed with either normal males or irradiated males. These results indicate that higher doses of gamma irradiation severely impair reproductive capacity, likely due to damage to reproductive tissues and genetic material (Puanmanee et al. 2010, Collins and Taylor 2011, Nasution et al. 2018). Zahran et al., (2013) also reported a decline in egg laying with increasing irradiation doses, with no eggs produced at doses of 50 Gy or higher when irradiated females were used. In contrast, Mahmoud and Barta (2011) found that gamma irradiation had no significant effect on egg production in *B. zonata* when irradiated males were mated with normal females.

Egg hatchability significantly declined with increasing radiation dose. When irradiated males were paired with normal females, hatchability was 33.45% at 30 Gy and decreased to10.64% at 70 Gy. . Complete (0%) hatchability was observed at 50, 60, and 70 Gy when irradiated females were crossed with either normal males or irradiated males. In contrast, the control group exhibited highest hatchability. Similar trends were reported by Panduranga et al. (2021) and Nasution et al. (2018). Nasution et al. (2018) found that egg hatchability declined significantly to 3.5% at 30 Gy, 0.37% at 50 Gy, and 0% at 70 Gy or higher.

Sterility reached 100% in irradiated females mated with either irradiated or normal males at doses of 50 Gy and above. While several studies have reported complete reproductive failure at different irradiation levels, the dose required to achieve dose full sterility may vary depending on factors such as dose rate, insect species, and the development stage of the pupae at the time of exposure (Burditt et al, 1975; Calkins et al, 1988). In our study, the highest sterility (97.4%) was observed when irradiated males were crossed with normal females at 70 Gy. This finding aligns with Calkins and Parker (2005), who reported that females are generally more radiosensitive than males. Similarly, Zahran et al. (2013) observed 100% sterility in irradiated females at doses of 50 Gy and above. When irradiated males were paired with normal females, the highest sterility rate (97.40%) was observed at 70 Gy.

This radiosensitivity may be attributed to the presence of nurse cells in female ovaries, which contain polytene chromosomes and large nuclei with unknotted chromatin resulting from endomitosis. . In contrast, male reproductive cells are more uniform and undergo fewer divisions, which may render them more resistant to radiation (Espo et al. 2015). Additionally, irradiation can disrupt spermatogenesis by damaging testicular development, reducing sperm motility, and inducing lethal mutations in sperm DNA (Kuswadi 2011, Espo et al. 2015).

Pupal recovery and adult emergence declined progressively with increasing irradiation doses. No pupae or adults emerged when irradiated females were crossed with either irradiated or normal males at doses of 50 Gy or higher. In contrast, some pupal recovery was observed at all irradiation levels when irradiated males were crossed with normal females. The lowest pupal recovery occurred at 70 Gy, supporting the findings of Embaby et al. (2022), who also reported significant reductions in pupation and adult emergence at higher radiation doses.

Pupal size, in terms of both weight and length, decreased with increasing irradiation dose, regardless of the mating combination. The smallest pupae were recorded at 70 Gy, consistent with the findings of Mahmoud and Barta (2011), who reported significant reductions in pupal size at this dose compared to the control.

Our results also revealed post-mating malformation in F1 progeny. A higher incidence of partial emergence and deformities was observed at 70 Gy in crosses involving irradiated males with un-irradiated females, as well as irradiated females with un-irradiated males. These findings suggest that radiation-induced damage can be transmitted to the offspring, leading to significant developmental disruptions. Such damage may impair the formation of internal organs, resulting in malformed adults with shrunken or undersized bodies and wrinkled wings. Ahmad et al. (2021) similarly reported heritable morphological abnormalities in F1 generations of irradiated *B. cucurbitae*, indicating transgenerational effects of genetic damage.

The present study provides baseline information on the effect of gamma irradiation on the biological and reproductive parameters of *B. zonata*, supporting its potential application in the Sterile Insect Technique (SIT). The findings indicate that irradiation doses of 60 and 70 Gy, applied to mature pupae (6 days old) using a Co^60^ gamma irradiator induced 94% and 97% sterility, respectively, in male fruit flies.

However, further studies are needed to optimize these irradiation doses to achieve complete sterility while preserving essential quality control traits such as male flight ability, mating competitiveness, and survival. Standardizing these parameters is crucial for the effective and sustainable implementation of SIT programs targeting *B. zonata*.

## Acknowledgements

The authors express their sincere gratitude to the Nuclear Institute for Food and Agriculture (NIFA), Peshawar, and the Institute of Zoological Sciences, University of Peshawar, for facilitating and supporting this research study. Special thanks are extended to Fazal-e-Rahim (PSA) at NIFA for his valuable assistance in laboratory management.

## Author contributions

Bibi Hajra (Conceptualization [equal], Writing—original draft [equal], Formal—analysis and data curation [equal], Methodology [equal], Project Management, Writing—review and editing [equal]), Muhammad Hamayoon Khan (Supervision, Investigation, Validation, Formal analysis [equal], Writing—review and editing [equal], Resources, Software [equal],), Farrah Zaidi (Supervision, Formal analysis [equal], Writing—review and editing [equal], Resources [equal]), Muhammad Salman (Writing—review and editing [equal]), Zainab Saeed (Writing—review and editing [equal]), Usman Khalique (Writing—review and editing [equal]), Mehreen Ayaz (Writing—review and editing [equal]), Syeda Hira Fatima (Writing—review and editing, [equal]), and Syed Jawad Ahmad Shah (Supervision, Writing—review and editing [equal], Resources [equal]).

